# Cholesterol sensing by CD81 is important for hepatitis C virus entry

**DOI:** 10.1101/542837

**Authors:** Machaela Palor, Lenka Stejskal, Piya Mandal, Annasara Lenman, Pia Maria Alberione, Jared Kirui, Rebecca Moeller, Stefan Ebner, Felix Meissner, Gisa Gerold, Adrian J. Shepherd, Joe Grove

**Author notes:** These authors contributed equally to this work (Machaela Palor, Lenka Stejskal).

## Abstract

CD81 plays a central role in a variety of physiological and pathological processes. Recent structural analysis of CD81 indicates that it contains an intramembrane cholesterol-binding pocket and that interaction with cholesterol may regulate a conformational opening of the large extracellular domain of CD81. Therefore, CD81 possesses a potential cholesterol sensing mechanism; however, the relevance of this for protein function is thus far unknown. In this study we investigate CD81 cholesterol sensing in the context of its activity as a receptor for hepatitis C virus (HCV). Structure-led mutagenesis of the cholesterol-binding pocket reduced CD81-cholesterol association, but had disparate effects on HCV entry, both reducing and enhancing CD81 receptor activity. We reasoned that this could be explained by alterations in the consequences of cholesterol binding. To investigate this further we performed molecular dynamic simulations of CD81 with and without cholesterol; this identified a potential allosteric mechanism by which cholesterol binding regulates the conformation of CD81. To test this, we designed further mutations to force CD81 into either the open (cholesterol unbound) or closed (cholesterol bound) conformation. The open mutant of CD81 exhibited reduced receptor activity whereas the closed mutant enhanced activity. These data are consistent with CD81 cholesterol sensing resulting in a switch between a receptor active and inactive state. CD81 interactome analysis also suggests that conformational switching may modulate the assembly of CD81-partner protein networks. This work furthers our understanding of the molecular mechanism of CD81 cholesterol sensing, how this relates to HCV entry and CD81’s function as a molecular scaffold; these insights are relevant to CD81’s varied roles in both health and disease.

## Introduction

Binding of the E2 glycoprotein of hepatitis C virus (HCV) to the large extracellular loop of CD81 is a defining event in the entry of HCV [1] and is targeted by multiple broadly neutralising antibodies, thus placing this molecular interaction at the forefront of current HCV vaccine development [2,3]. Whilst the importance of CD81 in HCV entry is well established, the precise details of E2-CD81 interaction have yet to be defined and the molecular determinants of CD81 receptor activity are only partially understood [4].

CD81 is a prototypical member of the tetraspanin superfamily. Tetraspanins are small integral membrane proteins, defined by their four transmembrane domains separated by intra/extracellular loops. Highly-conserved cysteine residues stabilise tetraspanin tertiary structure through disulphide bridges, and provide sites for post-translational palmitoylation, which influences tetraspanin membrane segregation [5,6].

Largely without cognate ligands, tetraspanins participate indirectly in a wide variety of cell-biological processes through their interactions with partner proteins, which they organise into functional complexes [7,8]. For example, CD81 facilitates the assembly of the B-cell receptor complex and is therefore essential for normal antibody responses. CD81 performs this role via partnership with CD19; first by chaperoning CD19 through the secretory pathway and then by dictating its cell surface distribution, permitting proper assembly of the B-cell receptor complex upon activation [9–12]. Through other molecular partnerships CD81 has been implicated in additional physiological processes such as T-cell receptor signalling, cell migration, growth factor signalling, sperm-egg fusion and most recently, biological ageing, potentially through its interaction with TMEM2 [13–19].

Aside from these physiological functions, CD81 is also commandeered by diverse infectious pathogens. It participates in the cell-surface assembly of both human immunodeficiency virus (HIV) and influenza A virus; a function that may be linked to the apparent affinity of CD81 for membrane structures with high curvature [20–23]. CD81 also negatively regulates SAMHD1 function, resulting in increased intracellular pools of dNTPs, which in turn favours HIV reverse transcription [24]. Finally, CD81 is critical for the entry of HCV and *Plasmodium* sporozoites into human hepatocytes [1,25]. In summary, CD81 performs molecular scaffolding function in a variety of pathways; a greater understanding of its molecular characteristics will provide novel insights into both physiological and pathological processes.

The recent crystal structure of CD81, the first of any tetraspanin, has provided a novel perspective on its molecular biology [26]. CD81’s four helical transmembrane domains are arranged in a loose bundle forming an inverted conical shape. Curiously, the transmembrane domains enclose a central intramembrane cavity filled by a single molecule of cholesterol, which is coordinated by hydrogen bonding to the side chains of inward-facing amino acids. Whilst this observation may have arisen due to the presence of cholesterol in the crystallisation buffer, Zimmerman et. al. use biochemical experiments to demonstrate physical association of CD81 with cholesterol [26]. Moreover, this finding i s consistent with other reports linking cholesterol to tetraspanin biology [27,28].

Whereas the minor extracellular domain (EC1) was not resolved in the crystal structure, CD81’s major extracellular domain (EC2) was found to be roughly parallel to the plane of the plasma membrane, analogous to a lid sitting on top of the bundle of transmembrane domains (Fig. 1A and Fig S2A). Overall, CD81 adopts a compact structure that is likely to project only a few nanometers from the cell surface. However, using molecular dynamic (MD) simulations Zimmerman et. al. demonstrated that the EC2 of CD81 has a propensity to flip up into an extended open conformation (Fig S2A). Furthermore, removal of cholesterol from the intramembrane cavity during the simulations increased the frequency of conformational switching, suggesting an allosteric link between cholesterol binding and CD81 conformation. These observations indicate that CD81 may have an, as yet unappreciated, function as a cholesterol sensor; this feature is likely to be important for its role as a scaffold for events occurring at cellular membranes.

**Fig. 1.**
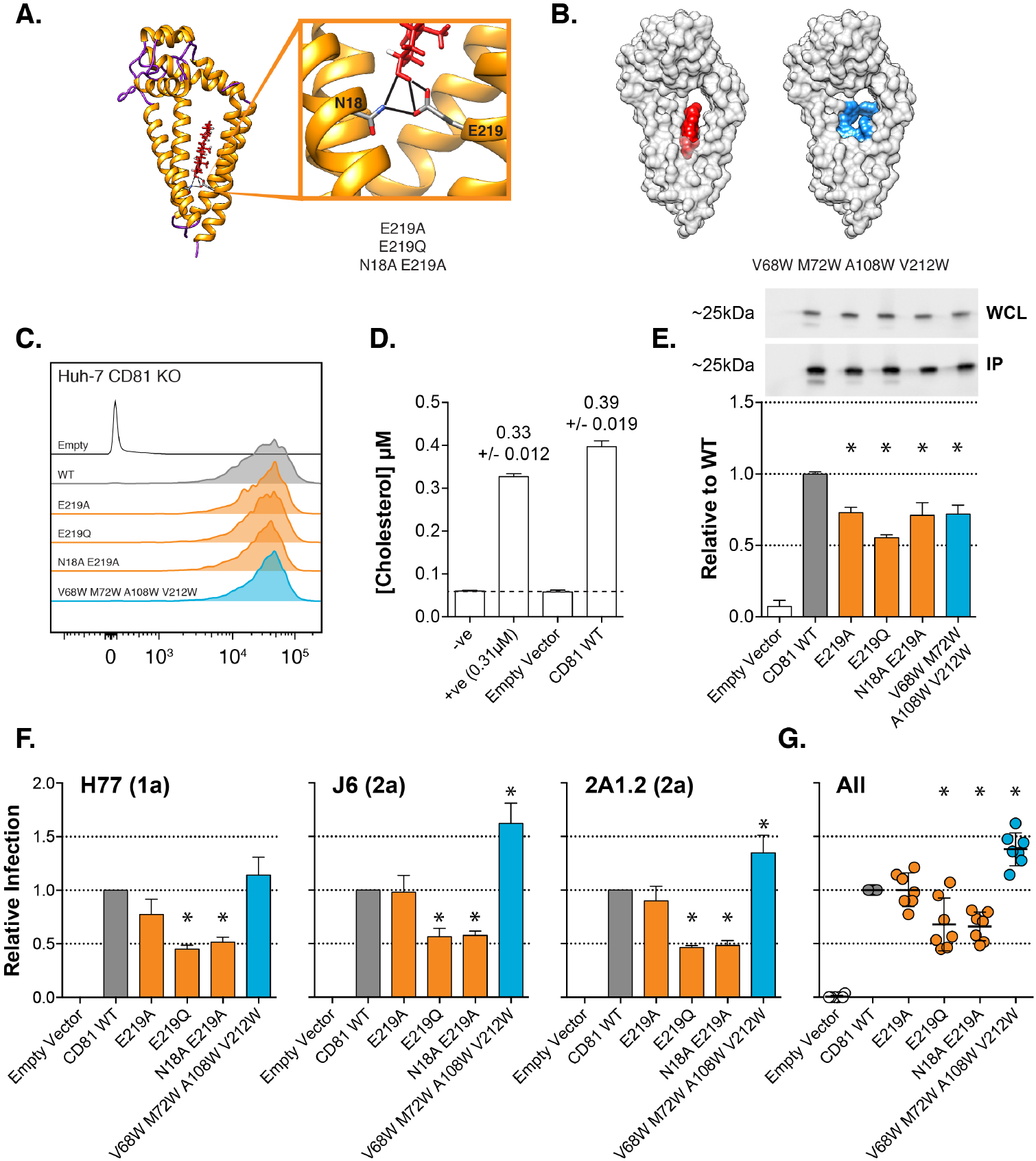
Mutations in the cholesterol binding pocket of CD81 modulate HCV entry. **A.** Cholesterol (red) is coordinated in the intramembrane cavity of CD81 by hydrogen bonds with inward facing residues N18 and E219. We made various mutations at these sites to disrupt this interaction. **B.** The cholesterol molecule sits in the centre of an intramembrane binding pocket. In the V68W M72W A108W V212W mutant this space is occupied by tryptophan residues (blue residues). **C.** The cell surface expression levels of each mutant CD81 was assessed by flow cytometry. **D.** Huh-7 CD81 KO cells were transduced with lentivector encoding WT CD81 or empty vector control. The cells were surface labelled with anti-CD81 mAb and lysed in Brij-98 detergent buffer. CD81-mAb complexes were pulled-down with protein G beads and associated free cholesterol was measured. Our positive control demonstrates the accuracy of the assay. The dashed line indicates the limit of detection **E.** We assessed cholesterol association with WT and mutant CD81. Data is expressed relative to WT CD81, asterisks indicate statistical significance from WT (n=4, one-way ANOVA, Prism). A representative western blot demonstrating equivalent levels of CD81 in the whole cell lysate (WCL) and pull-down (IP) **F.** Huh-7 CD81 KO cells were transduced with lentivectors encoding WT and mutant CD81. HCV entry was assessed by challenge with a panel of HCVpp (including genotypes 1, 2 and 5). HCVpp infection is shown relative to cells expressing WT CD81. Data from three representative clones and a summary plot of all HCVpp are shown. Asterisks indicate statistical significance from WT (n=4, one-way ANOVA, Prism). Error bars indicate standard error of the mean.

Whilst the precise molecular interaction of HCV E2 with the EC2 of CD81 has yet to be structurally defined, the relevant protein domains have been identified [29–34]. The CD81 binding site of HCV E2 comprises discontinuous protein regions, brought together in the 3D structure of the glycoprotein; these interact with helices D and E of CD81’s EC2, which are presented at the apex of CD81’s closed compact structure. Antibodies that prevent this interaction block HCV entry, and cells without CD81 are completely resistant to infection [35–44]. The ability of CD81 to recruit molecular partners is also likely to be important for HCV infection; indeed, other HCV entry factors constitutively associate with CD81 [8,45]. Significantly, HCV entry also seems to be closely linked to cell-surface cholesterol transport: three cholesterol-transporting proteins (SR-B1, LDLR and NPC1L1) have been implicated in the process [46]. Notably, the cholesterol transporter scavenger receptor B-1 (SR-B1) naturally associates with CD81 and also modulates the CD81-dependent invasion of *Plasmodium* sporozoites into hepatocytes [8,47,48].

The biology of both CD81 and HCV converge on plasma membrane cholesterol; therefore, we set out to investigate how CD81’s interaction with cholesterol impacts HCV infection. First, we mutated residues within the cholesterol binding pocket of CD81. Whilst all of the tested mutations reduced CD81-cholesterol association they had varying effects on HCV, both decreasing and increasing virus entry. This suggests the cholesterol binding pocket of CD81 is important for HCV infection, but viral entry may not be directly dependent on cholesterol association. We performed multiple independent molecular dynamics (MD) simulations of CD81 behaviour with and without cholesterol. In support of the report by Zimmerman et. al., we demonstrate a cholesterol-dependent conformational switch of CD81; this is consistent with the notion of cholesterol sensing by CD81. These experiments identified a potential hinging between CD81’s EC2 and transmembrane domains. We designed mutations to alter this motion and, therefore, disrupt CD81’s cholesterol sensing mechanism. Mutations that are predicted to confer the open conformation (i.e. the cholesterol unbound state) reduced HCV entry, whereas mutations that confer the closed (i.e. cholesterol bound state) enhanced HCV entry. Further characterisation of these mutants demonstrate that they exhibit normal cell surface expression and distribution, and retain the ability to chaperone CD19 to the cell surface. However, the open mutant of CD81 exhibits reduced interaction with HCV E2. We also use diverse cell culture proficient HCV to demonstrate that cholesterol-binding and open conformation mutants of CD81 do not support authentic HCV infection. This study provides further insight into the molecular mechanism of cholesterol sensing by CD81 and demonstrates that this activity is important for HCV infection.

## Results

### Mutations in the cholesterol-binding pocket of CD81 modulate HCV entry

The crystal structure of CD81 reveals an intramembrane cavity bounded by the four transmembrane domains; this contains a single molecule of cholesterol, which is coordinated by hydrogen bonding to the side chains of two residues, N18 and E219 (Fig. 1A). We designed a series of mutants to disrupt this interaction; E219A and E219Q, that were previously demonstrated to reduce CD81-cholesterol association [26], and an N18A E219A double mutation, which should remove all possibility of hydrogen bonding to cholesterol. Many of the inward-facing residues of CD81’s intramembrane cavity have small side chains (e.g. alanine, valine, glycine), this creates a binding pocket to accommodate cholesterol. Therefore, we also mutated four inward facing residues to tryptophan (V68W M72W A108W V212W), the side chain of which includes a bulky indole group. Structural modelling predicts that these tryptophan residues will fill the cholesterol-binding pocket, whilst maintaining the hydrophobic nature of the transmembrane domains (Fig. 1B). We introduced each of the cholesterol-binding pocket mutants into Huh-7 CD81 KO cells by lentiviral transduction and confirmed that their cell-surface expression was equivalent to WT CD81 (Fig. 1C).

Zimmerman et. al. previously demonstrated CD81-cholesterol association by the addition of exogenous cholesterol to purified CD81 [26]; we corroborated this by examining the interaction of CD81 with endogenous plasma-membrane-resident cholesterol. Huh-7 CD81 KO cells transduced with WT CD81 or empty vector control, surface labelled with anti-CD81 mAb and then lysed for immunoprecipitation. Following pull-down of CD81-mAb complexes with protein G beads we assayed the concentration of free, unesterified, cholesterol; this is the form of cholesterol found in cellular membranes [49]. Our negative control determined the limit of detection (~0.05*μ*M), whilst our positive control demonstrated the accuracy of the assay (Fig. 1D). In the pull-down from cells transduced with empty vector control we did not measure any free cholesterol, whereas cholesterol was readily detectable in the pull-down from cells expressing WT CD81.

Next, we went on to measure cholesterol association with the binding pocket mutants; each of the mutants exhibited a reduction in co-immunoprecipitated cholesterol (Fig. 1E). Notably, this experiment cannot discriminate between cholesterol that is directly associated with CD81 and peripheral cholesterol that is indirectly extracted during lysis and immunoprecipitation. Therefore, it is possible that the reduction in cholesterol concentration in each of our mutant pull downs represents a complete loss of specific cholesterol binding; this, however, cannot be determined. Nonetheless, these data are consistent with specific association between plasma-membrane cholesterol and CD81, and that this interaction can be reduced by mutating residues in the cholesterol-binding pocket of CD81.

We challenged Huh-7 cells expressing each mutant with a panel of HCV pseudoparticles (HCVpp); these are lentiviral reporters pseudotyped with the E1E2 glycoproteins of diverse strains of HCV, as such this system recapitulates the events of HCV entry [50,51]. Whilst the various cholesterol pocket mutants possessed identical cellular expression and similar deficiency in cholesterol association (Fig 1C & E) they exhibited differential HCV receptor activity (Fig 1F). Of the mutations that disrupt hydrogen bond formation with cholesterol, E219Q and N18A E219A reduced HCV entry by ~ 50%, whilst the E219A single mutant had equivalent receptor activity to WT CD81. Notably, the V68W M72W A108W V212W mutant, in which the binding pocket is filled with bulky tryptophan side chains, enhanced HCV entry by ~ 50%. These data demonstrate that mutations within the cholesterol-binding pocket of CD81 have the capacity to both negatively and positively modulate HCV entry. However, given that the level of cholesterol association did not correlate with receptor activity, it is unlikely that HCV is directly dependent on cholesterol occupying the intramembrane binding pocket of CD81.

### SR-B1 does not enhance cholesterol loading into CD81

CD81 constitutively associates with SR-B1 [7], a cell surface cholesterol transporting protein, which possesses a central hydrophobic tunnel through which cholesterol can be conveyed from high-density lipoproteins directly to the plasma membrane [52,53]. Moreover, this lipid transport function has been demonstrated to modulate the role of CD81 in HCV and malaria entry [48,54]. These observations led us to hypothesise that SR-B1 may directly load cholesterol into the binding cavity of CD81. Indeed, when human SR-B1 is over-expressed in CHO cells total cellular cholesterol levels double (Fig S1A). However, co-expression of SR-B1 did not alter CD81-associated cholesterol levels, as assessed by immunoprecipitation (Fig S1B). This is despite the detection of SR-B1-CD81 complexes in the pull down. The data presented in Fig S1B is from a single representative experiment; however, multiple repeat experiments in CHO and Huh-7 SR-B1 KO cells provided no evidence for cholesterol loading by SR-B1. Therefore, this hypothesis is not supported by the data.

### Cholesterol regulates conformational switching of CD81

Zimmerman et. al. reported that cholesterol binding regulates a switch in the EC2 of CD81 from a closed conformation (cholesterol bound) to an open conformation (cholesterol unbound) (Fig S2). This provides a molecular mechanism by which CD81 may sense cholesterol in cellular membranes. We reasoned that cholesterol sensing, rather than cholesterol binding in and of itself, may provide a mechanism by which the binding pocket mutants may modulate HCV entry. To investigate this further we used molecular dynamics (MD) simulation: an *in silico* methodology for predicting the conformational dynamics of proteins, both at steady state and after perturbations such as ligand removal or mutagenesis [55,56]. We conducted five independent 500ns MD simulations of CD81 with and without cholesterol and quantified the conformational state of the EC2. In the presence of cholesterol CD81 remained largely in a closed conformation, similar to that seen in the crystal structure, whereas in the absence of cholesterol the EC2 had a propensity to adopt a more extended open conformation. This was particularly apparent in a hinging motion between helix E of the EC2 and transmembrane domain 4 (TMD4) (Fig 2A). We therefore quantified the change in angle around this hinge in each simulation, using the angle adopted in the crystal structure as a reference (Fig 2B). In the absence of cholesterol, 3/5 simulations demonstrated clear and sustained extension of the hinge between the EC2 and TMD4. In the presence of cholesterol the angle of the hinge fluctuated in some simulations but provided little evidence of a persistent conformational switch. To further quantify this we calculated the cumulative time spent in the extended conformation across all five simulations, using an angle of 25° as a threshold (Fig 2C). This analysis suggests that in the absence of cholesterol CD81 is ~ 3 times more likely to be found in the open conformation.

**Fig. 2.**
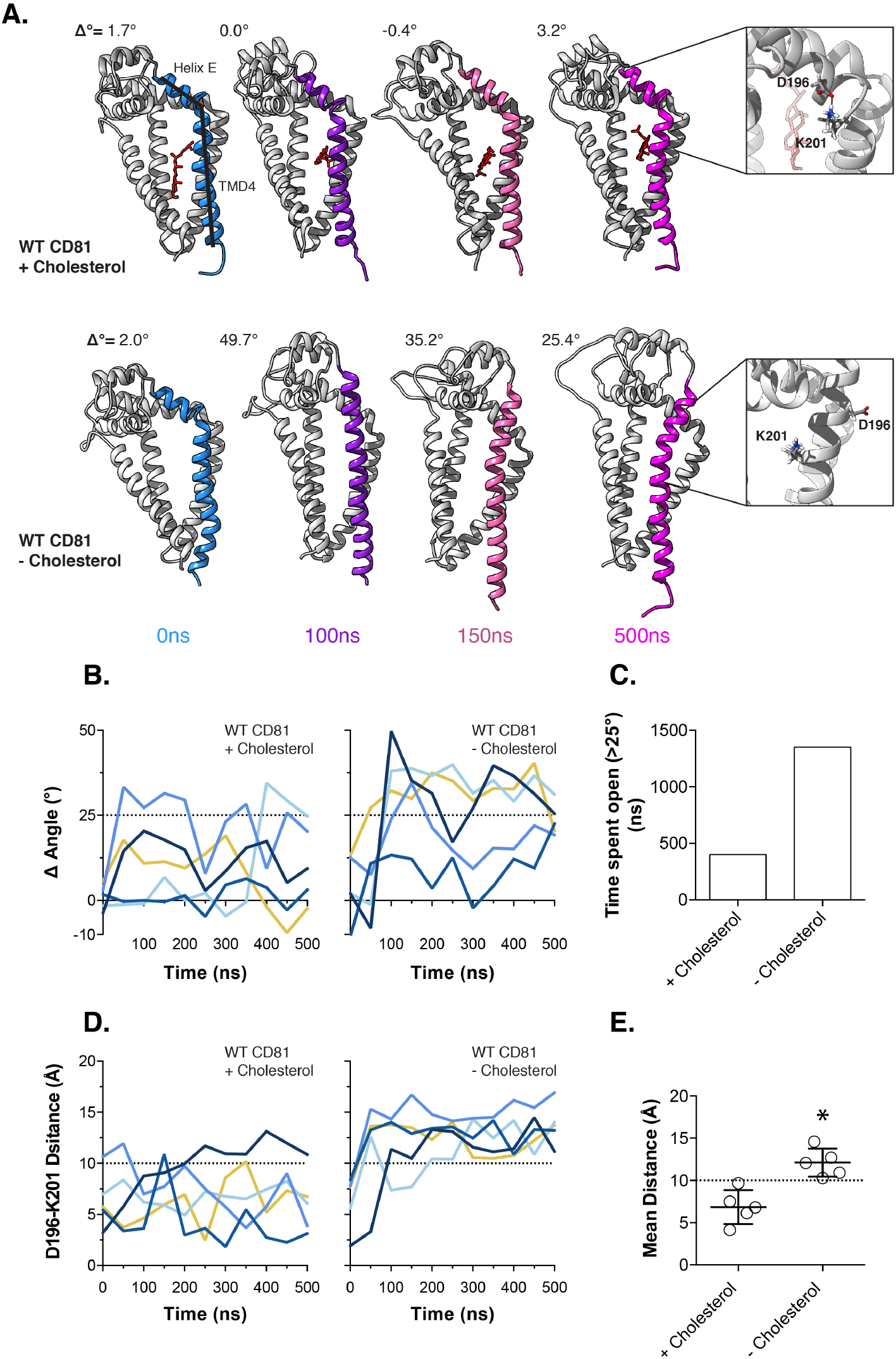
Conformational switching of CD81 in the absence of cholesterol. We performed five independent 500ns MD simulations of WT CD81 with and without cholesterol. **A.** Snapshots summarising representative simulations from either condition. The Δ° measurement reflects the change in the angle between helix E of the EC2 and TMD4 (as annotated), by comparison to the CD81 crystal structure. For each snapshot the region from which the measurement was taken is color-coded by time. Cholesterol is shown in red. Structures were orientated using TMD4 as a reference. Examples of the orientation of D196 and K201 are shown as insets. **B.** The angle between helix E and TMD4 was measured over time for each simulation, 25° was chosen as a threshold to indicate conformational switching. **C.** The cumulative time spent in the open conformation was calculated across all simulations for either experimental condition. **D.** The distance between D196 K201 was measured over time for each simulation, the dashed line indicates the distance under which electrostatic interactions and hydrogen bonding occurs (10Å). **E.** The average distance between D196 and K201 with and without cholesterol, data points represent the mean value for each simulation, asterisk indicates statistical significance (n=5 simulations, unpaired T-test, Prism).

In the cholesterol bound crystal structure of CD81 the hinge between the EC2 and TMD4 is stabilised in a closed conformation by a salt-bridge between D196 (in helix E of the EC2) and K201 (at the top of TMD4) (Fig S2B). Indeed, we observed interaction between these residues during our MD simulations in the presence of cholesterol (Fig 2A, top inset). However, without cholesterol CD81 adopts the open conformation, in which these residues are orientated on opposite sides of a continuous alpha-helix (Fig 2A, bottom inset). To examine this further we measured the distance between D196 and K201 during each simulation (Fig 2D). In the presence of cholesterol the distance between D196 and K201 fluctuates, nonetheless they remain in close proximity and frequently reach the short distances (<10Å) at which electrostatic interactions and hydrogen bonding can occur [57]. By contrast, in the absence of cholesterol these residues are consistently >10Å apart, indicating a change in orientation. To directly compare the experimental conditions we calculated the average distance between D196 and K201; we found the value to be significantly higher in simulations without cholesterol (Fig 2E).

Taken together, these data are consistent with the reports of Zimmerman et. al. and support the notion of a cholesterol-dependent conformational switch in CD81. Moreover, our MD simulations predict that absence of cholesterol disrupts stabilising interactions across the EC2-TMD4 hinge, presumably through allosteric reorientation of amino acid side chains. This relationship provides a potential molecular mechanism for cholesterol sensing by CD81.

### Conformational switch mutants modulate HCV entry

To investigate whether the conformational switch required for CD81 cholesterol sensing is relevant to HCV entry we mutated both D196 and K201 to alanine, therefore, preventing the possibility of interactions between these residues stabilising the EC2-TMD4 hinge. We would predict that the D196A K201A mutant would be more likely to adopt an open conformation, irrespective of the cholesterol binding status of CD81. To test this, we performed five independent MD simulations with the D196A K201A open mutant in the presence of cholesterol. In line with our expectations, the EC2-TMD4 hinge exhibited persistent opening in 2/5 simulations; consequently the D196A K201A was ~ 2 times more likely to be found in the open state (Fig 3 A, B & C). We introduced the D196A K201A open mutant into Huh-7 CD81 KO cells and challenged them with HCVpp; the N18A E219A cholesterol binding mutant was also included as a point of comparison (Fig 3E). D196A K201A exhibited poor receptor activity for all tested HCV strains and was statistically indistinguishable from the cholesterol binding mutant.

**Fig. 3.**
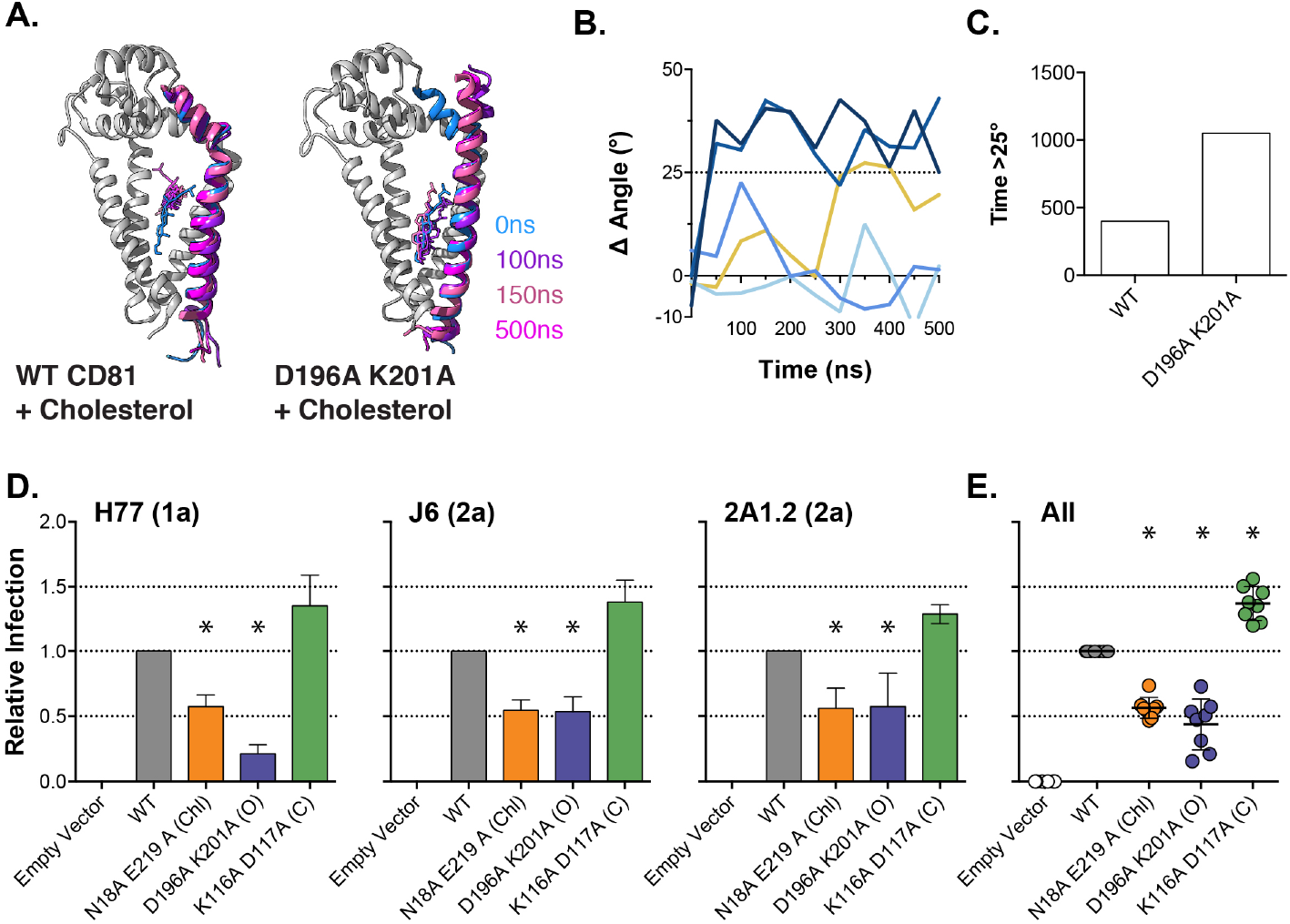
Conformational switch mutants modulate HCV entry. We mutated residues D196 and K201 to prevent stabilizing interactions across the EC2-TMD4 hinge. **A.** We performed five independent MD simulations of WT and D196A K201A CD81 in the presence of cholesterol. Images provide overlaid snapshots from representative simulations. Helix E, TMD4 and cholesterol are color coded by time. For clarity the remaining structure is shown in grey for the t=0ns snapshot only. Structures were orientated using TMD4 as a reference **B.** The change in angle between Helix E and TMD4, by comparison to the CD81 crystal structure, was measured over time for each D196A K201A simulation; compare to Fig 2B. **C.** The cumulative time spent in the open conformation for either WT or D196A K201A CD81. **D.** Huh-7 CD81 KO cells were transduced with lentivectors encoding WT CD81, N18A E219A (cholesterol binding mutant), D196A K201A (open mutant) or K116A D117A (closed mutant). HCV entry was assessed by challenge with a panel of HCVpp (including genotypes 1, 2, 4 and 5). HCVpp infection is shown relative to cells expressing WT CD81. Data from three representative clones and a summary plot of all HCVpp are shown. Asterisks indicate statistical significance from WT (n=4, one-way ANOVA, Prism). There was no significant difference between N18A E219A and D196A K201A. Error bars indicate standard error of the mean.

The open conformation of CD81 reported by Zimmerman et. al. (Fig S2A) is stabilised by a salt bridge between K116 and D117, which sits around a hinge between helix A of the EC2 and TMD3. We mutated these residues to destabilise the putative open state and, therefore, create a closed mutant of CD81 (Fig S2C); we tested this alongside the cholesterol binding and open mutants in the HCVpp assay. The K116A D117A mutant displayed a consistent enhancement of receptor activity (Fig 3D).

The fact the D196A K201A open mutant phenocopies the N18A E219A cholesterol mutant suggests that they both reduce CD81 receptor activity by inducing conformational opening. Whereas the opposite effect was observed for the K116A D117A mutant, which is expected to spend more time in the closed conformation. Therefore, these data support a model in which HCV entry is dependent on the cholesterol-mediated closed conformation of CD81.

### Residues that contribute to cholesterol sensing are highly conserved

CD81 is found in all vertebrates, Figure S3A displays the structure of CD81 colour coded for its amino acid conservation across vertebrates; the transmembrane domains and hinge regions of the EC2 display high conservation, whilst the apex of the EC2 exhibits increased diversity. This suggests that the outward-facing apex is under positive selection to drive new interactions and/or to escape pathogen binding [58], whilst the lower region of the protein remains conserved to maintain residues that are essential for basic protein function. If cholesterol sensing and conformational switching is an important feature of CD81 we may expect high conservation at the residues that regulate this process. Figure S3B is a phylogenetic tree constructed from representative vertebrate CD81 protein sequences and annotated to show the degree of conservation observed for the set of mutated residues described above. The residues found in the cholesterol-binding pocket and the conformation-switching residues are all conserved, suggesting functional importance. Notably, the residues at 116 & 117 and 196 & 201 maintain the potential for salt bridge formation in 15/15 and 13/15 of the representative sequences; salt bridges often stabilise conformational intermediates of a protein. As a point of comparison we include F186, which is presented at the apex of CD81 and is critical for HCV binding [34,59]; like much of the EC2, this position exhibits low conservation, consistent with the CD81-dependent species specificity of HCV infection.

### CD81 mutants retain normal trafficking a nd CD19 chaperone function

CD81 participates in diverse cell biological processes via its interactions with various binding partners [6]; for instance, CD81 is critical for B-cell receptor signalling by chaperoning CD19 through the secretory pathway [10–12]. Therefore, we evaluated the cellular distribution and chaperone function of the cholesterol sensing/conformational switch mutants to determine if this correlated with HCV receptor activity. In all subsequent experiments, as an additional control, we included an F186A mutant that is unable to bind HCV E2 and does not support virus entry [34,59].

When expressed in Huh-7 CD81 KO cells each of the mutants exhibit equivalent cell surface expression by flow cytometry (Fig 4Ai) and display no overt changes in cell surface distribution (Fig S4A). This suggests that the mutant’s receptor activities are not correlated with trafficking deficiencies. To assess chaperone function, we recapitulated CD81-dependent trafficking of CD19 in Huh-7 cells. Fig S4B displays exogenous CD19 expression in Huh-7 cells +/− CD81, as assessed by fluorescence microscopy; no cell surface CD19 is detectable in Huh-7 CD81 KO cells, despite equivalent total cellular expression of CD19 (Fig S4C). Fig 4Aii displays CD81-dependent cell surface expression of CD19 in Huh-7 cells. Each of the mutants maintains the ability to chaperone CD19 to the plasma membrane; these data indicate normal cell surface localisation for each of the mutants.

**Fig. 4.**
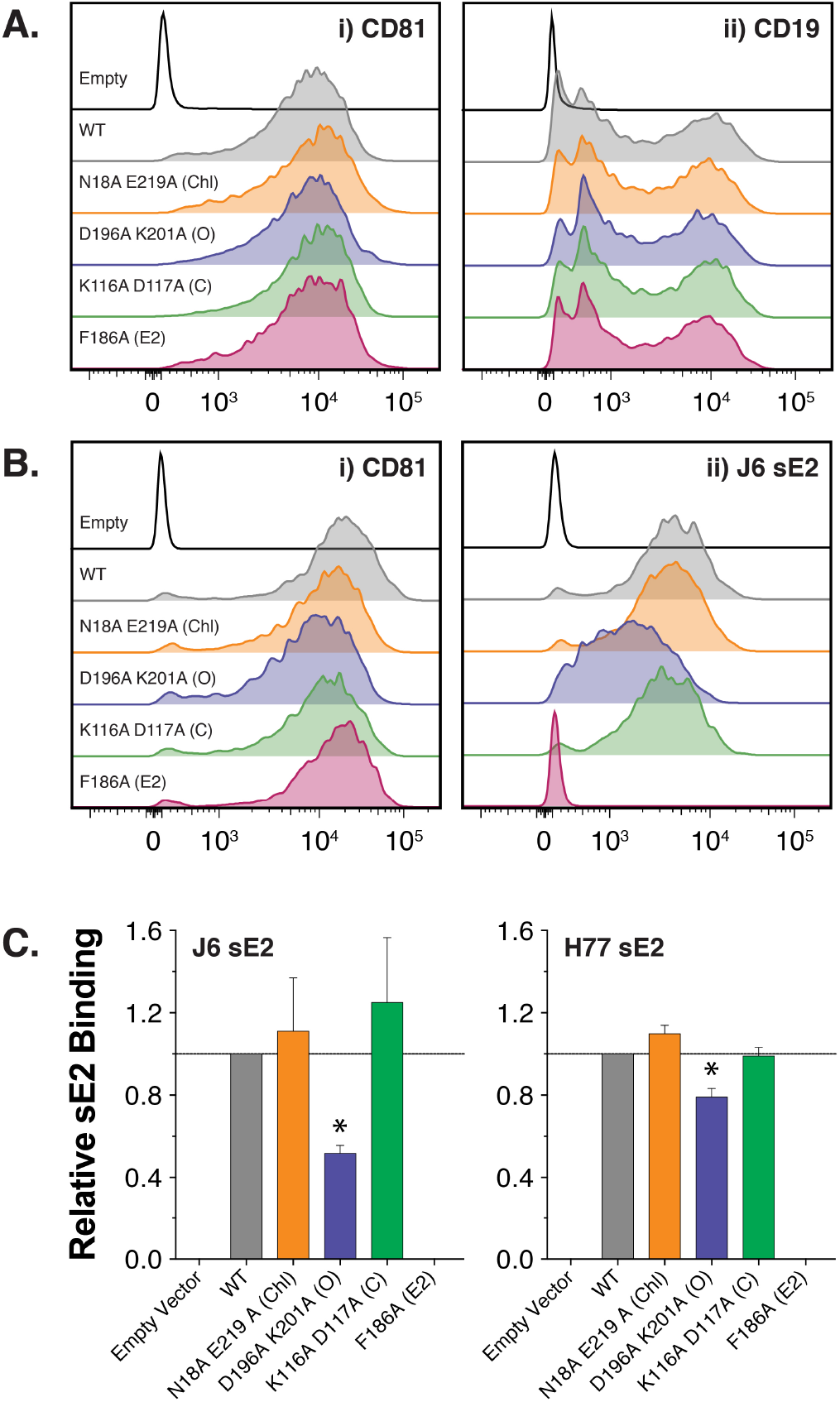
Cell surface functionality of CD81 mutants. Huh-7 CD81 KO cells were co-transduced with lentivectors encoding human CD19 and CD81 or empty vector. **A.** Representative flow cytometry histograms, all samples received CD19 lentivector plus the indicated CD81/control vector. The plot on the left demonstrates CD81 surface expression (i), the right-hand plot displays CD81-dependent trafficking of CD19 to the cell surface (ii). **B.** CD81 expression on CHO cells confers binding on soluble HCV E2. The plot on the left demonstrates CD81 surface expression (i), the right-hand plot displays sE2 binding to transduced CHO cells (ii). **C.** Quantification of sE2 binding expressed relative to WT CD81. Asterisks indicate statistical significance from WT (n=4, one-way ANOVA, Prism). Error bars indicate standard error of the mean.

CD81 facilitates HCV entry through interaction with the major viral glycoprotein E2. This can be examined experimentally using soluble E2 glycoprotein (sE2); whilst this is the least authentic system available to study HCV entry (sE2 is presented without its partner protein, E1, and not in the context of a virion) it is the only tool available to directly assess E2-CD81 interactions at the cell surface.

sE2 binding can be conferred to CHO cells by introduction of HCV receptors (CD81 or SR-B1) [60]. Therefore, we transduced CHO cells with the CD81 mutants and assessed binding of sE2 from the prototypical J6 and H77 strains. Figure 4Bi displays flow cytometry histograms of anti-CD81 mAb binding to each mutant, demonstrating equivalent CHO cell surface expression. Fig 4Bii provides representative plots of J6 sE2 binding to CHO-CD81 cells; note that whilst WT CD81 confers robust binding, the F186A mutant displays no interaction with E2. In this context, the N18A E219A cholesterol and K116A D117A closed mutants exhibit similar sE2 binding to that of WT CD81, whereas the D196A K201A open mutant has moderately reduced sE2 binding. This is quantified for both J6 and H77 sE2 in Fig 4C.

These data are not in particularly good agreement with the viral entry levels measured using HCVpp (Fig 3). Although the low receptor activity of D196A K201A is somewhat correlated with its ability to bind sE2, there is a disparity in effect sizes: e.g. D196A K201A exhibits only ~ 20% receptor activity for H77 HCVpp (Fig 3D) but retains ~ 80% binding to H77 sE2 (Fig 5C). Therefore, whilst it remains possible that differences in sE2 binding to CD81 mutants may contribute to their ability to support virus entry, it is likely that there is an additional determinant of receptor activity.

**Fig. 5.**
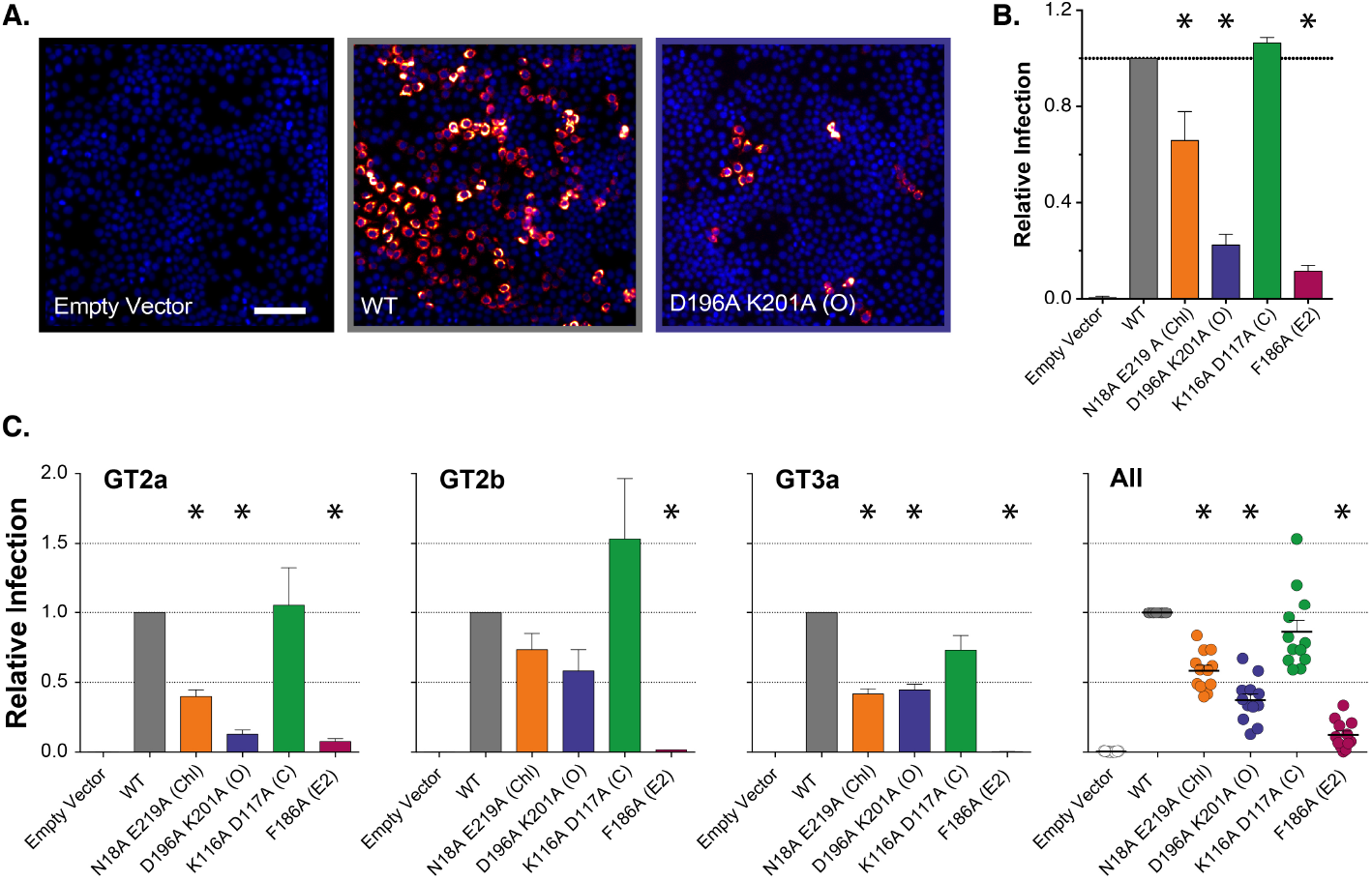
Cholesterol sensing is important for authentic HCV infection. Huh-7 CD81 KO cells were transduced with lentivectors expressing the stated CD81 mutants and were then challenged with J6/JFH HCVcc. **A.** Representative micrographs of HCVcc infection in transduced cells; DAPI nuclei shown in blue, viral antigen NS5A displayed in orange, scale bar = 100*μ* m. **B.** Quantification of infection, compiled from four independent experiments, data is expressed relative to infection in cells expressing WT CD81. **C.** Huh-7 Lunet N cells stably expressing the stated CD81 mutants were challenged with a panel of diverse HCVcc bearing the glycoproteins of genotypes 1, 2, 3, 4 and 5. Infection was quantified via a virally encoded luciferase reporter and is expressed relative to WT CD81. Data from three representative clones and a summary plot of all HCVcc are shown. Asterisks indicate statistical significance from WT (n=3, one-way ANOVA, Prism). Error bars indicate standard error of the mean.

### CD81 cholesterol sensing is important for infection by authentic HCV

Whilst the HCVpp and sE2 experimental systems allow the direct evaluation of HCV entry and receptor interaction, they present the viral glycoproteins in unphysiological contexts. Therefore, to complement and corroborate these experiments we assessed CD81 receptor activity using the HCV cell culture model (HCVcc); this generates native HCV particles, which have the unusual property of being enriched in lipids and associated with host lipoprotein components [36,61–64].

As before we transduced Huh-7 CD81 KO cells with each of our CD81 mutants and then challenged with J6/JFH HCVcc, quantifying the resultant infection by microscopy. Untransduced cells were completely resistant to HCVcc infection, whereas cells expressing WT CD81 were readily infected (Fig 5A). Much like the HCVpp system, the N18A E219A cholesterol-binding mutant and the D196A K201A open mutant display reduced ability to support HCVcc infection (Fig 5B). However, in this context the K116A D117A closed mutant failed to enhance infection, as we observed using HCVpp. The F186A E2 binding mutant exhibited minimal receptor activity. We also challenged CD81 expressing cells with a panel of HCVcc luciferase reporter viruses bearing the glycoproteins of diverse HCV strains from genotypes 1, 2, 3, 4 and 5. These data support our findings with J6/JFH HCVcc: N18A E219A and D196A K201A phenocopy each other, being poor receptors for HCV, whereas the K116A D117A mutant was equivalent to WT CD81. As an additional control we evaluated the effect of mutant CD81 on an HCV subgenomic replicon (SGR); this is a truncated HCV genome that lacks the structural proteins and, therefore, does not produce virus particles, but allows the processes of HCV translation and genome replication to be studied in isolation [65]. In this system the cells expressing CD81, whether WT or mutant, supported JFH-1 SGR replication at a similar level to cells without CD81; this is expected, given that CD81 is thought to act during virus entry. It also demonstrates that the CD81 phenotypes observed using the HCVcc systems cannot be attributed to HCV translation or replication, and must be a consequence of virus entry (Fig S5). In summary, mutants that alter CD81 cholesterol binding and/or conformational switching are poor receptors for HCVcc particle entry; this demonstrates that cholesterol sensing by CD81 is important for HCV infection.

### Conformational switch mutants have altered protein interaction networks

Like any tetraspanin, the activity of CD81 is defined by its molecular partnerships. Indeed, the interactions of CD81 with SR-B1, CLDN1 and EGFR are important for HCV entry, whilst CD81’s physiological functions are driven by various other partnerships with, for example, integrins (cell migration) and CD3 (T-cell regulation) [66,67]. The molecular mechanisms of tetraspanin-partner interactions remain poorly understood. To investigate the contribution of cholesterol binding and/or conformational switching to CD81’s interaction network we performed co-immunoprecipitation (co-IP) using an anti-CD81 EC2 mAb, followed by label-free quantitative (LFQ) mass spectrometry on cells expressing CD81 variants. Interaction partners were identified by statistical comparison of LFQ intensities from cells expressing CD81 to control cells without CD81. We identified multiple interacting proteins for WT CD81, consistent with our previous investigations of the CD81 interaction network, including well-described partners such as SR-B1 (SCARB1), claudin-1 (CLDN), epidermal growth factor receptor (EGFR), transferrin receptor (TFRC), calpain-5 (CAPN5), integrin*β*-1 (ITGB1) and CD151 (Figure 6A). Figure 6B displays enrichment values for these aforementioned proteins in co-IP/MS analyses of CD81 expressing versus control cells [7,8]. Intensity values were similar for most of these selected proteins, suggesting comparable interactions with WT and mutant CD81. Moreover, interactions with reported HCV entry factors (SR-B1, claudin-1, EGFR, transferrin receptor and calpain-5 [8,68–71]) were not correlated to receptor activity. For example, EGFR association was reduced for both the D196A K201A (open) and K116A D117A (closed) mutants (Figure 6A & Figure S6). Nonetheless, these mutants have opposing activities in the infectivity assays (Figure 3 & 5), suggesting that EGFR interaction is not a determinant of these phenotypes.

**Fig. 6.**
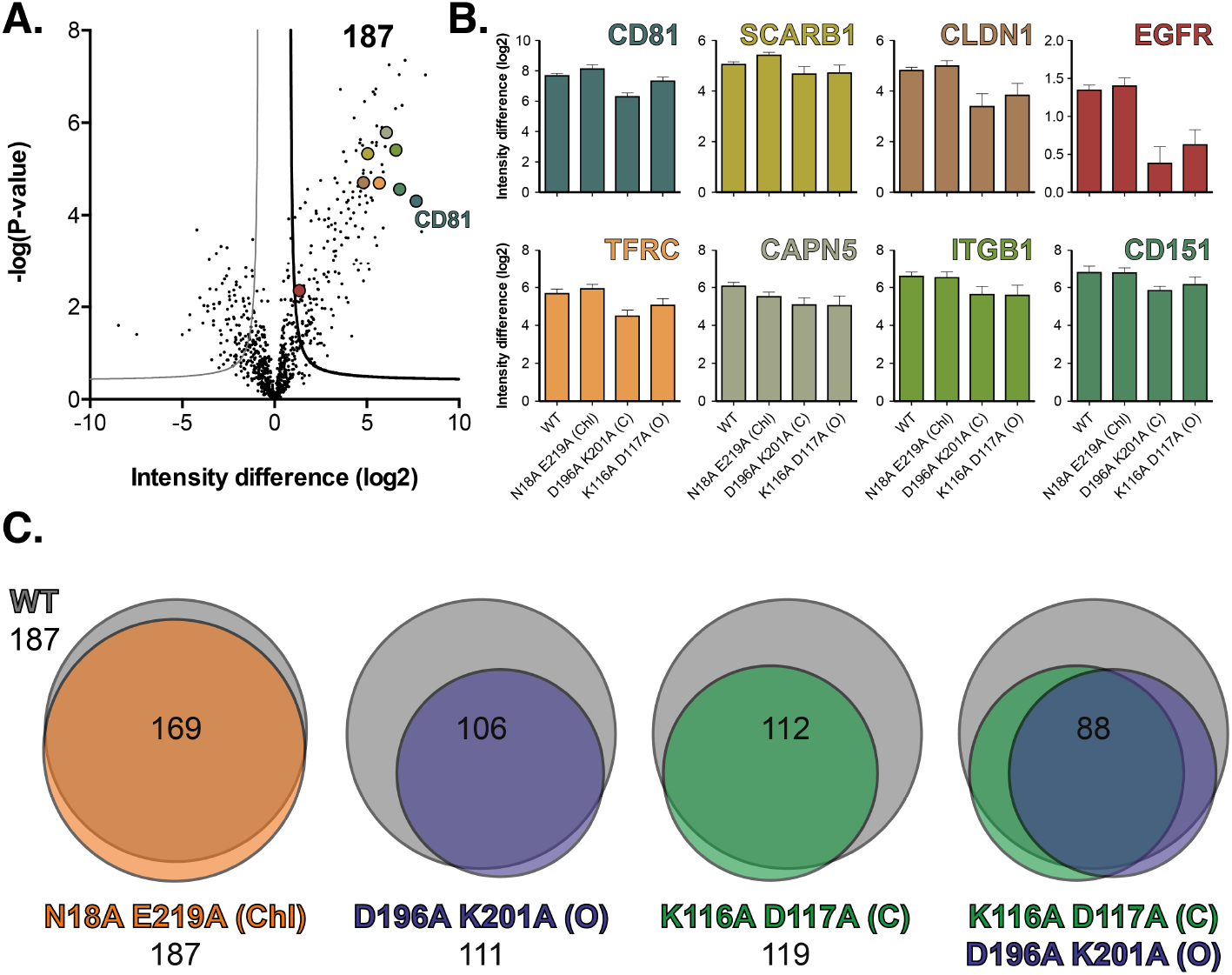
Conformational switch mutants exhibit altered protein interaction networks. **A.** Volcano plot visualizing differences from co-IPs of Huh-7 Lunet N CD81 WT versus Lunet N control cells (n=4 biological replicates for each cell line). LFQ intensity differences (log2) are plotted against the t-test p value (−logP). Significant interactors were defined by a permutation-based FDR using S0=1 as described [94]. Reference proteins (CD81, SCARB1, CLDN1, EGFR, TFRC, CAPN5 ITGB and CD151) are highlighted, color coded as in B. **B.** Mean LFQ intensity differences (log2) of interactors in CD81 co-IP (Huh-7 Lunet N CD81 WT and mutants versus Lunet N control cells). Error bars indicate standard error of the mean (n=4) **C.** Venn diagrams showing the overlap of significantly enriched proteins found in CD81 co-IPs from WT in grey, N18A E219A (Chl) in orange, D196A K201A (O) in purple and K116A D117A (C) in green. Values below each title indicate significant interactors for each CD81 variant, values in the center of each Venn diagram indicate overlapping interactors.

In addition to this focussed analysis of well-described interaction partners, we also compared the entire interaction network for each of the CD81 mutants. The cholesterol binding mutant (N18A E219A) was largely unchanged with >90% of interactions maintained. In contrast, the conformational switch mutants exhibit a 40-50% loss of interactions; cross comparison revealed a degree of polarisation between these mutants, suggesting the loss of different subsets of partnerships. These data suggest that the EC2 of CD81 is important for partner interactions, as has been recently demonstrated for CD19 [72]. Moreover, the conformation of the EC2 (either open or closed) may regulate protein network assembly. However, this experiment did not provide evidence of cholesterol-mediated regulation of partnerships, and we were unable to identify specific interaction(s) that correlated with HCV receptor activity.

## Discussion

Like many tetraspanins, CD81 partners with a variety of cell surface components to contribute to various physiological and pathological processes [5,6]. The CD81 crystal structure solved by Zimmerman et. al., provided a new perspective on tetraspanin biology [26] and proposed two novel features that may be highly relevant for protein function: i) CD81 is able to bind cholesterol in a cavity formed by its transmembrane domains and ii) the EC2 of CD81 undergoes a cholesterol-dependent conformational switch between a compact ‘closed’ form to an extended ‘open’ configuration. These properties bestow CD81 with a potential cholesterol-sensing mechanism; this is particularly interesting given the long-standing link between tetraspanin biology and cholesterol [7,27,28].

How cholesterol sensing contributes to CD81 function was, thus far, unclear. In this study, we investigated these structural features in the context of a well-characterised biological activity: CD81’s ability to mediate HCV entry. The importance of our work is two-fold; it provides a greater understanding of the molecular basis of CD81 cholesterol sensing, and also demonstrates that these features are important for CD81’s ability to support HCV infection.

First we mutated residues within the cholesterol-binding pocket of CD81 (Fig 1); these were designed to either directly prevent CD81-cholesterol interaction or to fill the binding pocket with bulky side-chains, therefore negating cholesterol binding. We demonstrated co-immunoprecipitation of CD81 and free cholesterol. This is good evidence of association between CD81 and plasma membrane-resident cholesterol. We observed a significant reduction in cholesterol association for each of our binding pocket mutants; this supports the notion of specific retention of cholesterol in CD81. However, cholesterol association did not reach background levels; this may indicate some non-specific association (due to detergent extraction) or that the mutations were insufficient to completely prevent capture of cholesterol.

Despite their equivalent deficiency in cholesterol association the binding pocket mutants had disparate activity in the HCVpp assay, being either neutral (E219A), detrimental (E219Q and N18A E219A), or beneficial (V68W M72W A108W V212W) to HCV entry. This suggests that receptor activity is not determined directly by the ability to bind cholesterol and may, instead, be the result of modulating the molecular consequences of cholesterol interaction. Interestingly, the enhancement of HCV entry by the V68W M72W A108W V212W mutant (and its phenotypic similarity to the K116A D117A closed mutant) may indicate that filling the binding pocket of CD81 with bulky side chains, in fact, mimics cholesterol binding.

Following these experiments we hypothesised that cholesterol-sensing, with its associated conformational change [26], may underpin the relationship between the cholesterol-binding pocket and HCV entry. To investigate this further we performed MD simulations of CD81 conformational dynamics; these experiments shed light on the potential molecular mechanisms of CD81 cholesterol sensing. In particular we observed a hinging motion between the EC2 and TMD4 of CD81 that was more likely to occur in the absence of cholesterol (Fig 2). Furthermore, in the CD81 crystal structure (which includes cholesterol) this hinge is conformationally locked in the closed orientation by a salt bridge between D196 and K201. By measuring the distance between D196 and K201 throughout the simulations we demonstrated that in the absence of cholesterol these residues are reorientated such that stabilising interactions are unlikely to occur. Therefore, we propose that cholesterol sensing occurs through allosteric regulation of the EC2-TMD4 hinge.

To test this potential mechanism we mutated the residues necessary for these stabilising interactions (Fig 3). The D196A K201A mutant performed as expected under MD simulation: conformational switching occurred more frequently than WT CD81 despite the presence of cholesterol in the binding pocket. The D196A K201A mutant also had poor receptor activity in the HCVpp system and was phenotypically indistinguishable from the N18A E219A cholesterol binding mutant. Furthermore, we designed an additional mutant (K116A D117A) that, based on the work of Zimmerman et. al. [26], is predicted to destabilise the open conformation of CD81; therefore, K116A D117A can be considered as being opposite to the D196A K201A. This closed mutant of CD81 exhibited enhanced HCV receptor activity and appeared to be phenotypically similar to the mutant with the filled cholesterol pocket (V68W M72W A108W V212W).

The results of these experiments can be reconciled with a potential model of CD81 cholesterol sensing and HCV infection. In its cholesterol bound state CD81 is in a closed conformation, it is this form of CD81 that is active for HCV entry. Conversely, conformational opening of CD81 in the absence of cholesterol reduces its receptor activity. Therefore, binding pocket mutants may modulate HCV entry by dysregulating cholesterol sensing: the E219Q and N18A E219A mutants reduce HCV entry by reducing cholesterol retention, whereas the V68W M72W A108W V212W enhances entry by mimicking cholesterol occupancy of the binding pocket.

In this framework, the open and closed mutants act by de-coupling conformational switching from cholesterol binding. The E219A mutant was neutral in the HCVpp assay, this is less consistent with the above model. However, our work suggests that cholesterol binding regulates CD81 conformation through an allosteric reorientation of the EC2-TMD4 hinge. Therefore, it is possible that the E219A mutant has a compensatory effect that prevents allosteric reorientation in the absence of cholesterol and, therefore, retains HCV receptor activity. Each of the mutated residues exhibit very high levels of conservation (Fig S3), which is consistent with their being necessary for an essential function, such as cholesterol sensing.

We used the HCVcc system to test the mutants in a more physiologically relevant setting. The results of these experiments were largely consistent with the HCVpp entry assay (Fig 5) with one notable exception: the K116A D117A closed mutant did not enhance HCVcc infection. This disparity is most likely attributable to the fundamental differences between the constituents of HCVpp and HCVcc particles, the latter being associated with host lipoprotein components and enriched for host-derived lipids, including cholesterol [62–64,73]. This may suggest that the lipoprotein-like properties of native HCV particles allow them to modulate CD81 cholesterol sensing.

CD81 participates in a variety of cell biological processes through its interactions with myriad binding partners. We used co-IP and LFQ mass spectrometry to characterise the interaction networks of the cholesterol sensing mutants (Fig 6). The conformational switch mutants (both open and closed) exhibited reduced interactions. This is consistent with the notion that the EC2 is responsible for binding partnerships and CD81 oligomerization [72,74,75]; moreover these data may suggest that EC2 conformation regulates protein network assembly. However, mutations to reduce cholesterol binding (N18A E219A) had a negligible effect on the interactome of CD81. This may indicate that cholesterol binding/sensing is inconsequential for CD81 interactions, although it is also possible that residual retention of cholesterol (Fig 1) is sufficient to allow normal partnership interactions. This relationship, therefore, requires further investigation.

We did not, however, determine why HCV entry may favour the closed conformation of CD81. We found no differences in the trafficking or cell surface distribution of mutant CD81, only the D196A K201A mutant exhibited any alteration in interaction with sE2 (Fig 4), and the interaction networks of the various mutants were not predictive of HCV receptor activity (Fig 6). Nonetheless, one might speculate on alternative mechanisms by which EC2 conformation determines receptor activity. For instance the close physical proximity of the closed EC2 to the plasma membrane may be beneficial. A recent report of the structure of HIV-1 gp120 in complex with its co-receptor CCR5 suggests that the principal role of CCR5 is to anchor the HIV fusion machinery in close apposition to the host membrane [63]; consequently the HIV fusion peptide is within striking distance upon activation. CD81 has been implicated in HCV fusion [64] and has a low profile at the cell surface, particularly in the closed conformation, therefore, it could perform an analogous anchoring function for the HCV glycoproteins. Under this mechanism, the open conformation of CD81 may generate an insurmountable gap between the viral and host membranes, inhibiting fusion and, therefore, entry.

Notably, SR-B1 funnels cholesterol from its lipoprotein ligands directly into the plasma membrane [52,76]. Moreover, it is well established that HCV entry is enhanced by lipoproteins and that HCV particles themselves are lipoprotein-like being highly enriched for cholesterol [62–64,73,77]. Therefore, we hypothesised that local delivery of cholesterol via SR-B1, may regulate CD81 function. However, co-over-expression of SR-B1 did not enhance cholesterol association with CD81 (Fig S1); this was observed in CHO cells and Huh-7 SR-B1 KO cells. Therefore, our data do not support this hypothesis.

Finally, it is worth considering how cholesterol sensing may function in the context of CD81’s natural cellular environment. The lipid milieu within distinct subcellular membranes is tightly regulated to impart different biophysical and functional properties [78]. For example, whereas the plasma membrane is enriched for free cholesterol, intracellular membranes (e.g. endoplasmic reticulum and Golgi apparatus) have 5-10 fold lower cholesterol content. Therefore, newly synthesised CD81 travels along a cholesterol concentration gradient as it passes through the secretory pathway to the plasma membrane; consequently cholesterol sensing is likely to occur most frequently at the cell surface. This provides a potential mechanism for regulating CD81 function in a location-dependent manner. Moreover, the cell surface does not present a homogenous lipid environment; phase separation results in microdomains enriched for different lipid species, including cholesterol [49]. Cell surface proteins segregate within these lipid microdomains and this can alter protein function [79,80]. It is possible that cholesterol sensing determines CD81 function in a microdomain-specific manner; indeed, there is evidence that CD81 can transition in and out of cholesterol-dependent lipid domains at the cell surface [28,47]. Therefore, cholesterol sensing may allow CD81 to respond to dynamic changes in the local lipid environment; how this might affect CD81’s physiological and pathological activities will be the focus of future research efforts.

## Materials and methods

### Cell culture

Huh-7 CD81 KO cells were a kind gift from Prof. Yoshiharu Matsuura (Osaka University, Japan) [37]. Huh-7.5 cells were acquired from Apath LLC. Huh-7 Lunet N cells were generated as previously described [81]. HEK293T and CHO-K1 cells were acquired from the American Type Culture Collection. All cells were grown at 37°C in DMEM supplemented with 10% foetal calf serum, 1% non-essential amino acids and 1% penicillin/ streptomycin.

### Antibodies

Anti-NS5 and anti-CD81 mAbs were a kind gift from Prof. Jane McKeating (University of Oxford, UK). The anti-CD81 mAbs have been described in detail recently [35]. Anti-CD19 (sc-19650) was purchased from Santa Cruz Biotechnology. StrepMAb classic was purchased from IBA GmbH (Göttingen, Germany). All secondary antibodies purchased from Thermo Fisher Scientific (Waltham, MA, USA).

### Lentiviral vectors

Commercially synthesised gene sequences encoding WT and mutant CD81 were inserted into lentiviral expression plasmids by restriction digest. CD19 was cloned into the same background through PCR amplification from human cDNA. These plasmids will be made freely available after publication: https://www.addgene.org/Joe_Grove/. To generate lentiviral vectors HEK293T cells were co-transfected with pCMV-dR8.91 packaging construct, pMD2.G VSV-G expression plasmid and one of each of the CD81 encoding plasmids. Supernatants containing viral vectors were collected at 48 and 72 hours. The transduction efficiency of vectors were titrated by flow cytometry to allow equivalent transduction and expression of CD81 variants.

### HCV pseudoparticles

HCVpp were generated in a similar manner to the lentiviral expression vectors. HEK293T cells were cotransfected with pCMV-dR8.91 packaging construct, a luciferase reporter plasmid and an expression vector encoding the appropriate HCV glycoprotein. Supernatants containing HCVpp were collected at 48 and 72 hours. UKN4.1.1, 5.2.1, 2A1.2 and 2B1.1 E1E2 expression plasmids were kindly provided by Alex Tarr and Jonathan Ball (University of Nottingham, UK), all other E1E2 plasmids were generated in-house through PCR or commercial gene synthesis.

### Cell culture proficient HCV

HCVcc were generated as described previously [36]. Briefly, in vitro transcribed full-length HCV RNA genomes were electroporated into Huh-7.5 cells. Supernatants containing infectious HCVcc were harvested every 2-4 hours during the day, from 3-7 days post electroporation. Harvests were then pooled, aliquoted and frozen to generate a standardised stock for infection assays.

### Infections

Huh-7 or Huh-7 Lunet N cells were seeded into 96 well plates 24 hours prior to the experiment; to infect they were challenged with HCVpp/HCVcc supernatants (diluted 1/2 – 1/4 in DMEM 6% FCS). The infections were allowed to proceed for 72 hours before read out. For HCVpp, the samples were lysed and assayed using the SteadyGlo reagent kit and a GloMax luminometer (Promega, Maddison, WI, USA). To measure HCVcc replication, cells were fixed with 100% methanol and stained for viral NS5 protein, the proportion of infected cells was determined using the ImageJ Infection Counter plugin [82], these data were also verified by manually counting foci forming units. For HCVcc encoding a Renilla luciferase reporter, cells were lysed with milli-Q water and frozen at −80 ° C to ensure complete lysis, the samples were transferred to 96 wells-white plates and mixed with Renilla luciferase substrate solution (Coelenterazine, 0.42 mg/ml in Methanol) and luciferase activity was determined using a microplate reader Centro XS (Berthold Technologies, Harpenden, UK).

### Subgenomic replicon system

SGR transcripts were generated by in vitro transcription and introduced into target cells by electroporation (as with HCVcc). Replication was assessed at 4 and 24 hours post electroporation by read out of a genetically encoded Renilla luciferase reporter, as described above.

### Flow cytometry

To measure cell surface expression of CD81 or CD19, single-cell suspensions of Huh-7/CHO cells were fixed in 1% formaldehyde and then blocked in PBS + 1% BSA. All subsequent steps are performed in blocking buffer. Cells (100μl at 1-3×10^6^/ml) were then serially incubated with anti-receptor antibodies followed by anti-mouse Alexa Fluor 647 secondary, 1 hour incubation each at room temperature. Fluorescence signals were measured on a LSR Fortessa (BD, Franklin Lakes, NJ, USA) and data was analysed using FlowJo (FlowJo LLC, Ashland, OR, USA). The lentiviral vectors, described above, also express GFP (from a separate promoter); therefore, this GFP signal was used as an independent measure of transduction to identify positive cells during analysis.

### Immunoprecipitation and cholesterol association assay

Huh-7 CD81 KO cells were transduced to express various CD81 cholesterol-binding mutants. To perform the pull-down, a confluent T150cm^2^ flask of cells were trypsinized, harvested and resuspended in ‘traffic stop’ buffer, PBS + 1% bovine serum albumin (BSA) and 0.01% sodium azide; this depletes cellular ATP pools, consequently preventing receptor internalisation [60]. The resuspended cells were then incubated with anti-CD81 mAb 2.131 at 1*μ*g/ml for 60 minutes. Cells were washed with PBS and then lysed in 1% Brij58 buffer (1% Brij58, 20mM Tris pH 7.5, 150mM NaCl, 1mM CaCl_2_, 1mM MgCl_2_ and 0.02% NaN_3_) plus 1X protease inhibitor for 30 minutes. All steps were carried out on ice. Lysates were centrifuged at 13,300 rpm for 12 minutes at 4°C and then incubated with protein G sepharose beads for 90 minutes on a tube rotator in the cold room. Finally, beads were washed and stored at −20°C in 1% Brij58 buffer until further analysis.

We quantified cholesterol associated with CD81 using an Amplex Red Cholesterol Assay (Thermo Fisher Scientific, Waltham, MA, USA). Beads were pelleted by centrifugation, resuspended in 1X reaction buffer (as described by the manufacturer) and incubated in a tube rotator at room temperature for 15 minutes, this step extracts cholesterol associated with the immunocomplexes. Beads were then centrifuged and the supernatant was diluted 1/2 in 1X reaction buffer to a total volume of 50*μ*l. Samples were incubated with 50*μ*l working solution of Amplex Red Reagent at 37°C; cholesterol esterase was omitted from the CD81 samples to ensure the measurement of free-cholesterol only. Signal was measured using the fluorescence plate reading capability of a real-time PCR machine (Bio Rad, Hercules, CA, USA). A standard curve was prepared through serial dilution of the cholesterol reference in 1X reaction buffer, allowing the concentration of cholesterol to be determined in the positive control and CD81 samples.

### Western Blot

One day prior to study, Huh-7 cells +/− CD81 and/or CD19 were seeded into standard 24 well plates at 4×10^4^ cells/well. Cells were then lysed using a buffer containing 20mM Tris-HCl, 135mM NaCl, 1% Triton-X 100 and 10% glycerol. The samples were then run on a TruPage 4-12% gel under non-reducing conditions and transferred on to nitrocellulose membrane. The blots were blocked in PBS + 2% milk solution + 0.1% Tween-20 and then probed by serial incubation with anti-receptor antibodies and goat anti-mouse secondary conjugated to horseradish peroxidase. Chemiluminescence signal was then measured in a Chemidoc MP (Bio Rad, Hercules, CA, USA).

### Microscopy

One day prior to study, Huh-7 cells +/− CD81 and/or CD19 were seeded into standard 24 well plates at 1.2×10^4^ cells/well. Cells were then fixed (in s itu) i n 2% formaldehyde, blocked and stained, as described for flow cytometry, with the inclusion of a 10 minute DAPI counterstain at the end of the procedure. Samples were imaged on a Nikon Ti inverted microscope, through a 40X extra-long working distance objective, using a C2 confocal scan head with 405nm and 635nm laser illumination (Nikon Instruments, Tokyo, Japan). Multiple Z-stacks were acquired for each sample. Data was processed for display using FIJI/ImageJ [83,84].

### Soluble E2 binding assay

J6 and H77 E2 ectodomains (residues 384-661) were PCR cloned into expression vectors, as previously described [60], with the resultant constructs including an upstream tissue plasminogen activator signal sequence (to direct efficient secretion) and a downstream strep-tag II (for detection and purification). P roteins w ere produced by transient transfection of HEK293T cells with the supernatants being harvested at 48 and 72 hours post infection. sE2 was purified using a Strep-Tactin column (IBA Life Sciencens, Göttingen, Germany), monomeric sE2 was subsequently isolated by size exclusion chromatography. The sE2 binding assay was performed as previously described. A single-cell suspension of CHO +/− CD81 cells were preincubated in ‘traffic stop’ buffer, described above. All subsequent steps are performed in traffic stop buffer. Cells (100*μ* l at 1-3×10^6^/ml) were then mixed 10*μ*g/ml sE2 and incubated for 1 hour at 37°C. Bound sE2 was then detected using 3*μ*g/ml StrepMab classic followed by an anti-mouse Alexa Fluor 647 secondary, 1 hour incubation each at room temperature. Finally, cells were fixed in 1 % formaldehyde. Fluorescence signals were measured by flow cytometry.

### Co-immunoprecipitation and LC-MS/MS analysis

To identify CD81 interaction partners, Huh-7 Lunet N cells expressing CD81 (WT or mutant) or empty vector control were harvested from a 90% confluent T 150cm^2^ p late by scraping. Experiments were conducted in four biological replicates from four independent cell passages. Cells were lysed in Brij58 buffer (1% Brij 58, 50 mM Hepes, pH 7.4, 150 mM NaCl, 10 % glycerol, 1 mM CaCl_2_) supplemented with 1x protease and phosphatase inhibitors for 30 min on ice. Nuclear debris was pelleted at 12000g for 10 minutes at 4°C followed by co-immunoprecipitation using Pierce Crosslink Immunoprecipitation kit (Thermo Fisher Scientific, Waltham, MA, USA) with crosslinked anti-CD81 (clone 1.3.3.22, Santa Cruz). Efficiency of bait enrichment was determined by western blot using anti-CD81 (JS-81, BD Biosciences).

The co-IP samples were reduced, alkylated and trypsinized as previously described [7]. Peptide mixtures were analysed using a nanoflow liquid chromatography (LC-MS/MS) on an EASY-nLC 1000 system (Thermo Fisher Scientific) coupled to a Q Exactive HF-X Quadrupole-Orbitrap mass spectrometer (Thermo Fisher Scientific). A column oven (Sonation) maintained the temperature at 60° C. 500ng peptide samples in buffer A (0.1% formic acid) were loaded onto a 50-cm column with 75-*μ*m inner diameter, packed with C18 1.9 *μ*m ReproSil beads (Dr Maisch GmbH). Peptides were separated chromatographically with a 95 min gradient from 5% to 30% buffer B (80% acetonitrile, 0.1% formic acid).

The MS was operated in data-dependent acquisition mode, where one full scan (300 to 1650 m/z, R = 60000 at 200 m/z) at a target of 3×10^6^ ions is followed by 15 data-dependent MS/MS scans with higher energy collisional dissociation [target 10^5^ ions, max ion fill time 28 ms, isolation window 1.4 m/z, normalized collision energy 27%, R = 15000 at 200 m/z]. Dynamic exclusion of 30 s was enabled.

MS raw files were processed in MaxQuant (Version 1.6.2.0; [85]) and peptide fragment lists were searched against the human FASTA Uniprot reference proteome (June 2019) by the built-in Andromeda search engine. We restricted enzyme specificity with cleavage C-terminal after K or R (trypsin and Lys-C), allowing up to two missed cleavages. Fixed modifications of cysteine carbamidomethylation and variable modifications for the N-acetylation of proteins and the oxidation of methionine were specified. The minimum peptide length was set to seven amino acids. False discovery rates (FDRs) at the peptide and protein levels were 1%. The label-free-quantification (MaxLFQ) and matching between runs features were enabled.

Protein groups were filtered for decoys, contaminants and modifications. Data was also filtered for valid values (3, in at least one group). Interaction profiles were analysed with imputation of missing values (Width: 0.3, down shift: 1.8). Statistical analysis of proteomics data was conducted using two-sample t-test comparing LFQ intensities of proteins found in CD81 WT or mutants, against empty vector control (FDR: 0.05, s0=1). The resulting protein interactions have been submitted to the IMEx (http://www.imexconsortium.org) consortium through IntAct [86] with the assigned identifier IM-28053. The mass spectrometry proteomics data have been deposited to the ProteomeXchange Consortium via the PRIDE partner repository [87] with the dataset identifier PXD019260.

### Molecular dynamics simulations

We started with a molecular model of full-length CD81 in a closed conformation; this is based on the crystal structure (5TCX), as previously described, and was generously provided by Prof. Ron O. Dror and Dr. Brendan Kelly. All substitutions were introduced using Modeller software and AutoSub.py script available at https://github.com/williamdlees/AmberUtils. Protonation states were determined in MolProbity[88]. There are three histidine residues (37, 151, 191) within the CD81. In the models of WT without cholesterol and D196A K201A with cholesterol, all histidines were protonated on the epsilon nitrogen. In the WT with cholesterol model, the histidines 37 and 191 were also protonated on the epsilon nitrogen and the histidine 151 was protonated on the delta nitrogen. Models were then inserted into simulated palmitoyl-oleoyl-phosphatidylcholine (POPC) bilayers using CHARMM-GUI [88,89].

For MD simulations, each variant model was put through the same pipeline. First, the models were solvated in a rectangular box using TIP3P water molecules and 0.15 M of NaCl. The volume of the box was ~ 5 × 10^5^ Å^3^ with the total of ~ 5.2 × 10^4^ atoms including around 128 lipid molecules. The CHARMM 36 force field was used for the simulations on GPUs using the CUDA version of PMEMD in Amber 18[90–92]. The systems were minimised by 2500 steps of steepest descent followed by 2500 steps of the conjugate gradient method with all protein atoms restrained by a force of 10 kcal/mol/Å^2^ and phosphate atoms of the POPC bi-layer by a force of 2.5 kcal/mol/Å^2^. The systems were then heated to 310K for 25 ps using the Langevin thermostat under constant volume whilst keeping the identical restraints, followed by further 25 ps with the protein atoms restrained by 5 kcal/mol/Å^2^ and the phosphate atoms of the POPC bi-layer by a force of 2.5 kcal/mol/Å^2^. Initial velocities were sampled from Boltzmann distribution.

Further equilibration was performed under constant pressure (1bar) using the Monte Carlo barostat and semi-isotropic pressure coupling for 25 ps with the protein atoms restrained by 2.5 kcal/mol/Å^2^ and the phosphate atoms of the POPC bilayer by a force of 1 kcal/mol/Å^2^. This was followed by additional three equilibration steps lasting 100 ps each under constant pressure using the Monte Carlo barostat and semi-isotropic pressure coupling with the protein atoms decreasing to 1, 0.5 and 0.1 kcal/mol/Å^2^ and the phosphate atom restraints decreasing to 0.5, 0.1 and 0 restraints kcal/mol/Å^2^ respectively.

Following minimisation and equilibration steps, 500ns production runs were simulated under constant pressure using the Monte Carlo barostat, semi-isotropic pressure coupling and constant temperature via the Langevin thermostat at 310 K. For each independent simulation the restart file from the final equilibration step was used as the input for a short (1ns) production run but only the coordinates, not the velocity, were used to de-correlate the simulation. The coordinates from this short run were then used as input for the 500ns production run.

1 fs time step was used for minimisation, and the first two equilibration steps. SHAKE was used in order to restrain hydrogen bonds in all but the minimisation steps and 2 fs time-step was used for the last four equilibration and production runs. For all simulations, the cutoff distance for Lennard-Jones (LJ) 6-12 interactions was set to be 12 Å. The LJ 6-12 were smoothed over the range of 10 to 12 Å using the force-based switching function. Particle mesh Ewals (PME) method was used for the long-range electrostatic interactions and 1-4 nonbonded interactions were not scaled. To avoid the overflow of coordinates, the *iwrap* was set to 1.

### Sequence conservation analysis

Vertebrate CD81 encoding gene sequences were pulled from the NCBI database. Multiple sequence alignment and phylogenetic tree construction (using representative gene sequences) was performed using CLC sequence viewer (Qiagen, Hilden, Germany).

### Molecular modeling

Molecular graphics and analyses performed with UCSF Chimera, developed by the Resource for Biocomputing, Visualization, and Informatics at the University of California, San Francisco, with support from NIH P41-GM103311 [72].

### Statistical analysis

All statistical analysis was performed in GraphPad Prism 6.0 (San Diego, USA). Ordinary one-way ANOVA was performed using Dunnett’s multiple comparison test, using WT CD81 as a control, unless stated otherwise. Unpaired t-test was performed assuming equal standard deviation using a two-tailed p-value.

## Supporting information

Compiled Supplement

## ACKNOWLEDGEMENTS

We are grateful to Prof. Jane McKeating for reagents and advice. Thank you to Prof. Greg Towers and Mphatso Kalemera for scientific criticism. Thanks to Dr. Michael Tomlinson for technical advice on immunoprecipitation of tetraspanins. Also, thank you to Nicole Finardi for her efforts on the project. JG is supported by a Sir Henry Dale Fellowship from the Wellcome Trust and Royal Society (107653/Z/15/Z). LS received a Wellcome Trust PhD studentship (109162/Z/15/Z). GG was funded by the Knut and Alice Wallenberg Foundation, the Deutsche Forschungsgemeinschaft (DFG, German Research Foundation) - Projektnummer 158989968 - SFB 900 project C7 and DFG project GE 2145/3-2; the German Academic Exchange Service (DAAD) to JK; the Master Infection Biology, Alemania, Argentina (AMIBA) and CUAA/DAHZ to PA; and the Infection Biology International PhD Program of Hannover Biomedical Research School to RM.

## AUTHOR CONTRIBUTIONS

This study was conceived, designed and supervised by LS, FM, GG, AJS and JG. Biochemical and infection assays were performed by MP, PM, PMA, JK and RM. Computational biology was performed by LS. AL and SE performed mass spectrometry and bioinformatics analysis. FM advised in experimental design and analysis of mass spectrometry experiments. The manuscript was written by JG.

## Notes

### Competing Interest Statement

The authors have declared no competing interest.

### Summary of Updates

Extensive revision following peer review and further experiments since original version. Minor typo in author's name corrected since previous version.

