## Supplementary material for "Cholesterol sensing by CD81 is important for hepatitis C virus entry": Compiled Supplement

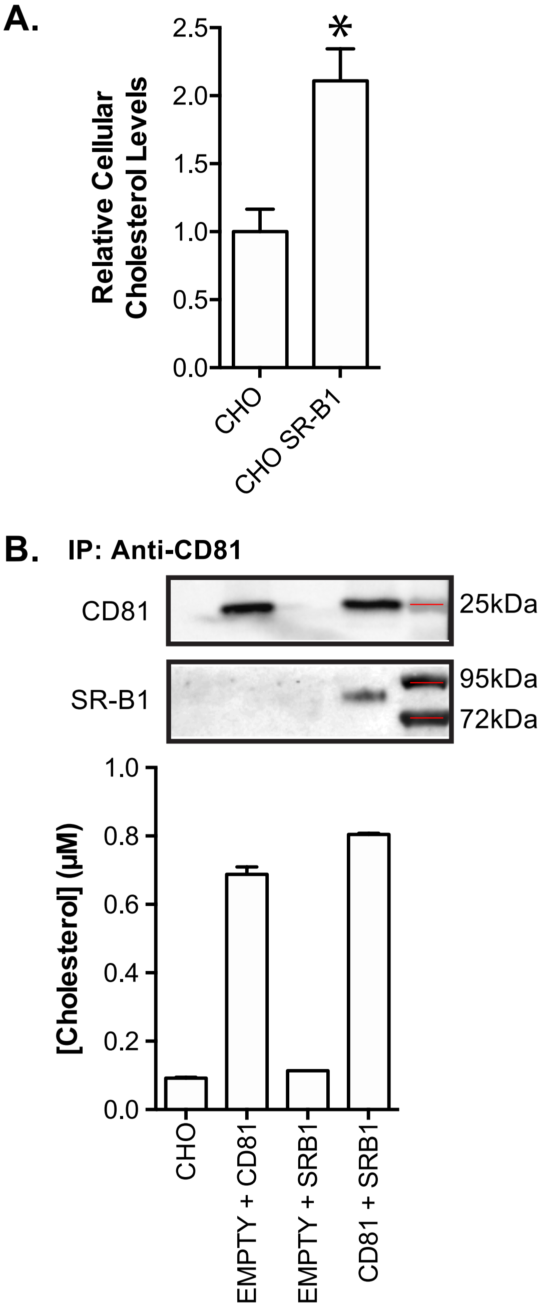


**Figure S1 SR-B1 does not enhance cholesterol loading into CD81. A.** Total cholesterol was measured in untreated CHO cells, or those transduced to express human SR-B1. Cholesterol levels were normalised to those in untreated cells. Values represent the mean of n=6 independent measurements, asterisk indicates statistical significance (T-test, Prism). Error bars indicate standard error of the mean. **B.** CHO cells were transduced with combinations of lentivector encoding human CD81 and/or SR-B1 and/or empty vector control. The cells were surface labelled with anti-CD81 mAb and lysed in Brij-98 detergent buffer. CD81-mAb complexes were pulled-down with protein G beads and associated protein and cholesterol was measured. The western blot displays protein detected after anti-CD81 immunoprecipitation, molecular weight markers are denoted by red lines. The plot, below, displays the cholesterol concentration in each sample. Error bars indicate standard error of the mean. Data is from one representative experiment.

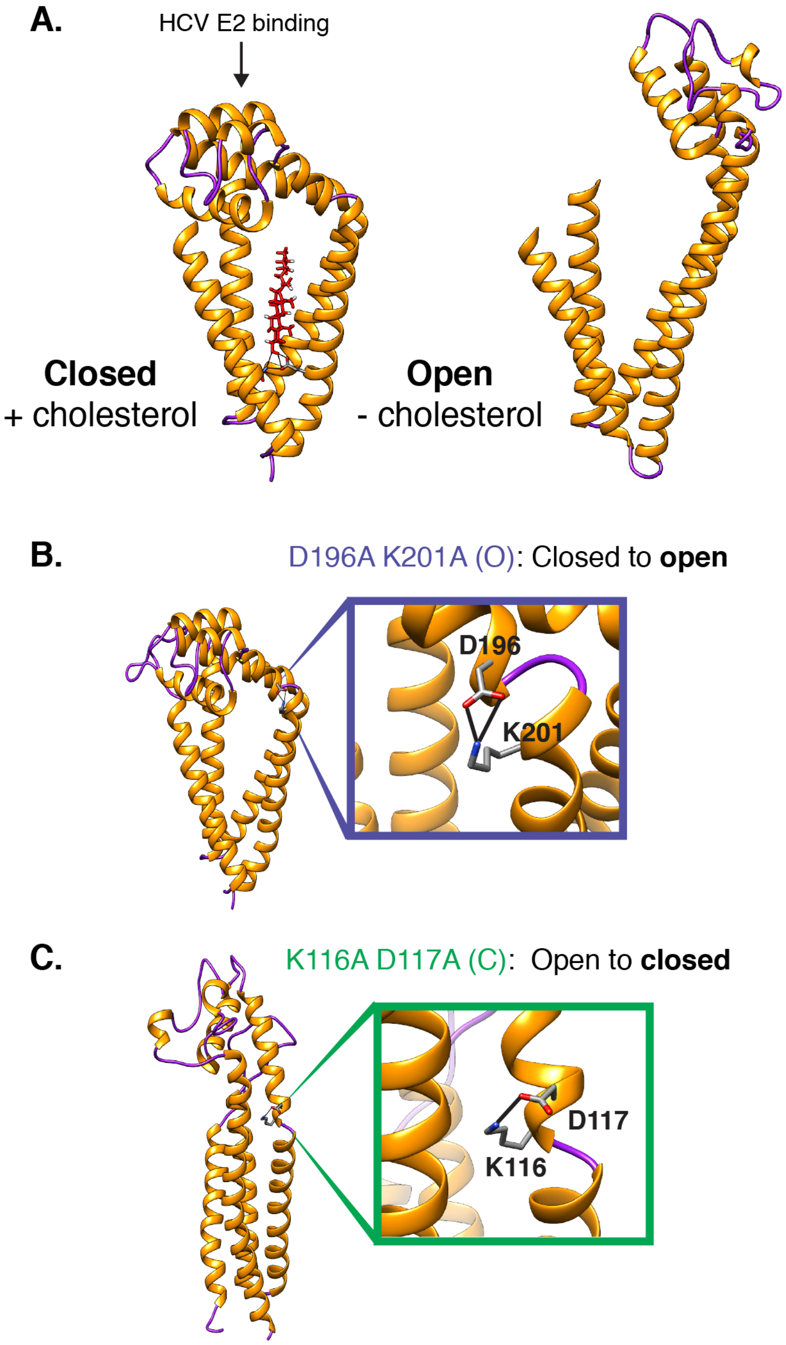


**Figure S2 Conformational switching of CD81 A.** Molecular models of CD81 in the closed (+ cholesterol) and open (- cholesterol) conformations; the closed conformation is based on the crystal structure (PDB: 5TCX), whereas the open conformation was generated by molecular dynamic simulations, as described by Zimmerman et.al [26]. Alpha helices are shown in gold, whereas unstructured regions are in purple, cholesterol is shown in red. HCV binds to helices D and E of the EC2, as indicated. The small EC1 is unstructured in both the open and closed conformations and has been omitted from the image for clarity. **B.** The closed conformation of CD81 is stabilised by a salt bridge between D196 and K201, which sit on the hinge described in Fig 2; we mutated these residues to alanine to create an open conformation mutant of CD81 (i.e. the conformational state in the absence of cholesterol). **C.** The work of Zimmerman et. al. predicts that the open conformation of CD81 is stabilised by a salt bridge between K116 and D117; we mutated these residues to alanine to create a closed mutant of CD81 (i.e. the cholesterol bound state).

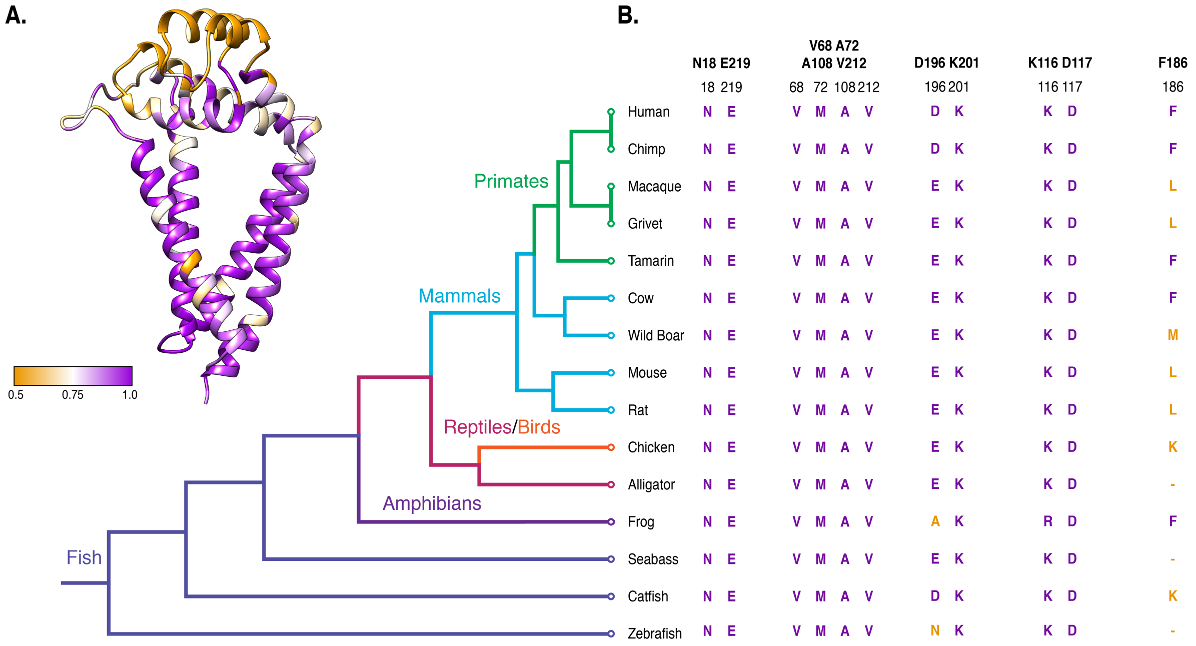


**Figure S3 Residues important for cholesterol sensing are highly conserved. A.** CD81 model (based on the crystal structure) with each residue colour-coded for conservation, purple indicates high conservation whilst gold represents low conservation (<0.5), as annotated. The conservation score was generated from an alignment containing 228 vertebrate CD81 genes. **B.** A phylogenetic tree containing representative vertebrate species; the amino acid identity at each site targeted by mutation is annotated. The letters are colour-coded, similar to A., to indicate conservation or divergence, relative to the sequence of human CD81.

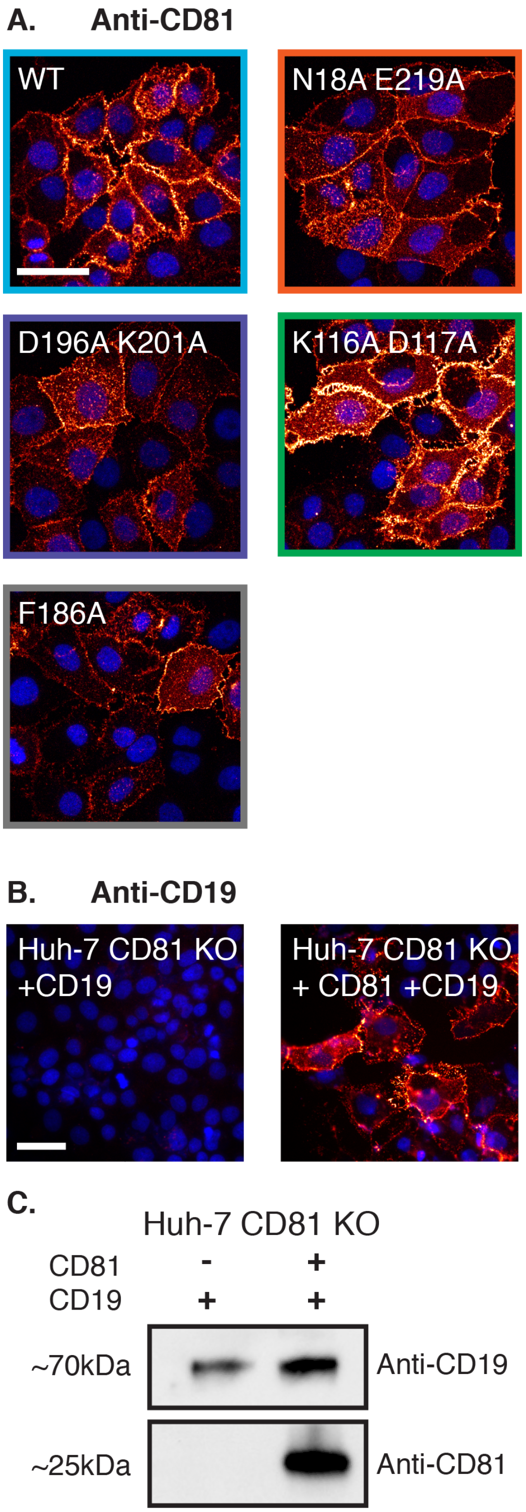


**Figure S4 CD81 mutants exhibit normal cell surface distribution.** Huh-7 CD81 KO cells were transduced to express CD81 variants and then fixed for microscopy. **A.** Representative maximum-intensity projections of cells stained with anti-CD81 2.131 (orange) and DAPI (blue), scale bar 50µm. **B.** CD81-dependent trafficking of CD19. Representative micrographs of cells transduced to express CD19 +/- CD81. Samples were stained with anti-CD19 (orange) and DAPI (blue). **C.** Parallel cultures were processed for western blot analysis. Representative images of CD19 and CD81 expression in transduced cells.

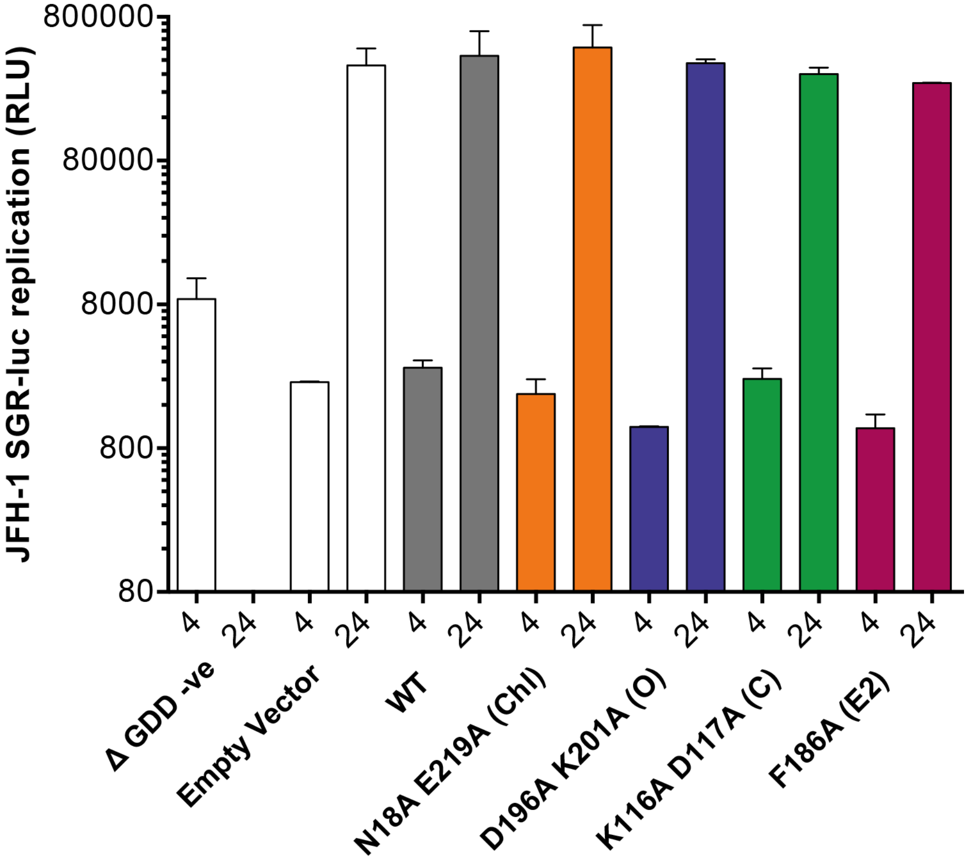


**Figure S5 A HCV subgenomic replicon is unaffected by mutant CD81 expression.** Huh-7 Lunet N cells stably expressing the stated CD81 mutants were electroporated with JFH-1 SGR or JFH-1 ΔGDD SGR replication incompetent negative control. Luciferase reporter activity was measured at 4 and 24 hours post electroporation: the signal at 4 hours represents translation from input RNA, the signal at 24 hours is a consequence of active replication. As expected, JFH-1 ΔGDD showed no evidence of active replication when introduced into cells expressing WT CD81; this was the case for each cell line (data not shown). JFH-1 SGR exhibited robust replication at 24 hours in each cell line, irrespective of mutant CD81 expression.

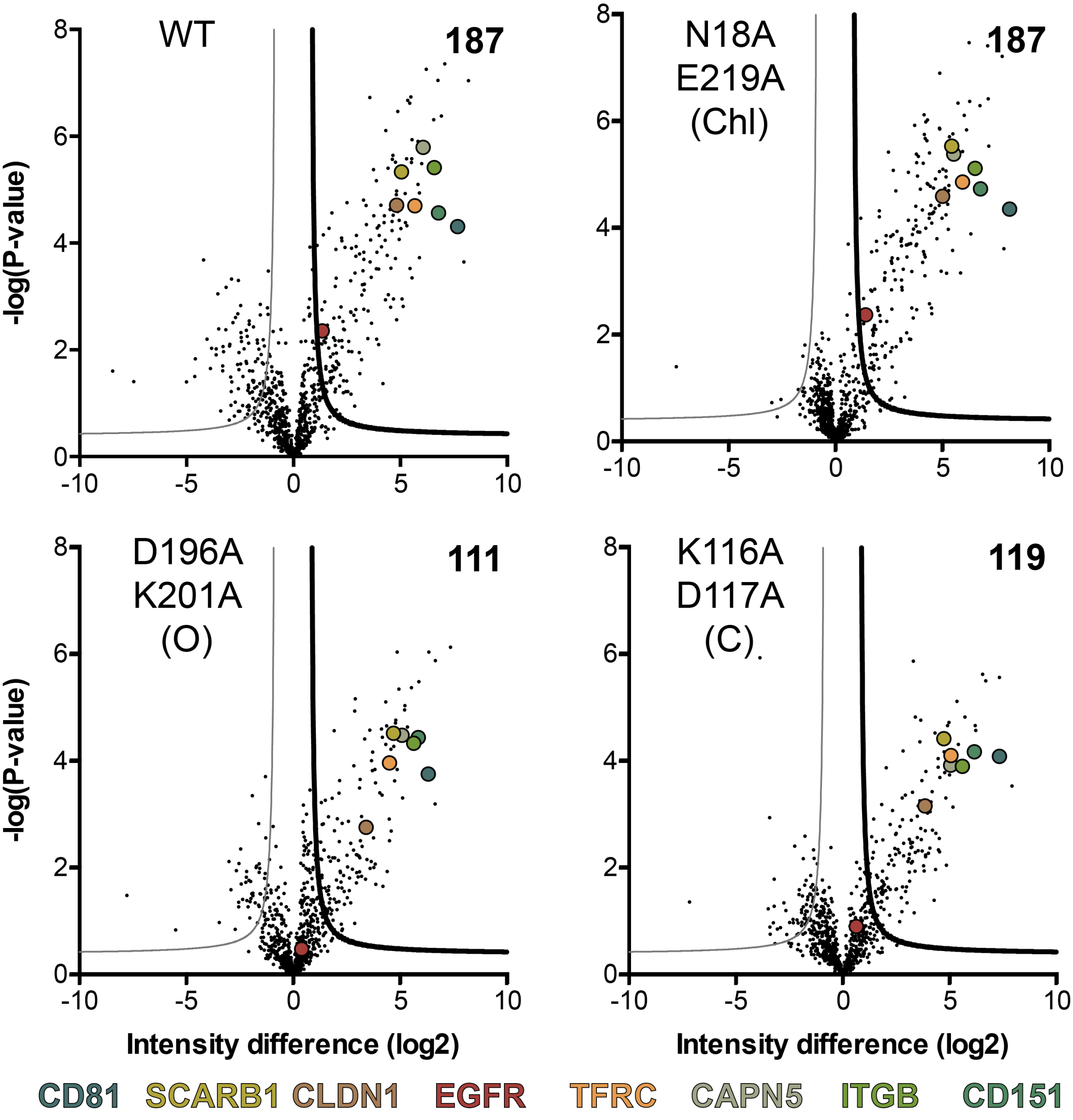


**Figure S6 LFQ mass spectrometry of CD81 WT and mutants. A.** Volcano plot visualizing differences from co-IPs of Huh-7 Lunet N CD81 WT/mutants versus Lunet N control cells (n=4 biological replicates for each cell line). LFQ intensity differences (log2) are plotted against the t-test p value (-logP). Significant interactors were defined by a permutation-based FDR using S0=1 as described for Figure 6. Reference proteins (CD81, SCARB1, CLDN1, EGFR, TFRC, CAPN5 ITGB and CD151) are highlighted as indicated in the color key.
